# High throughput, error corrected Nanopore single cell transcriptome sequencing

**DOI:** 10.1101/831495

**Authors:** Kevin Lebrigand, Virginie Magnone, Pascal Barbry, Rainer Waldmann

## Abstract

Droplet-based high throughput single cell isolation techniques tremendously boosted the throughput of single cell transcriptome profiling experiments. However, those approaches only allow analysis of one extremity of the transcript after short read sequencing. We introduce an approach that combines Oxford Nanopore sequencing with unique molecular identifiers to obtain error corrected full length sequence information with the 10×Genomics single cell isolation system. This allows to examine differential RNA splicing and RNA editing at a single cell level.

---

Single cell RNA sequencing (scRNA-seq) is a key technique for the analysis of cell-to-cell heterogeneity and projects aiming at analyzing the transcriptome of all cells from complex organisms have been initiated (e.g. Human Cell Atlas ^1^, Tabula Muris ^2^). While droplet based high throughput scRNA-seq approaches (e.g. 10×Genomics Chromium) allow the analysis of thousands of cells, they only yield limited sequence information close to one extremity of the transcript after Illumina short read sequencing. Information crucial for an in-depth understanding of cell-to-cell heterogeneity on splicing, chimeric transcripts and sequence diversity (SNPs, RNA editing, imprinting) is lacking.

Long read sequencing can overcome this limitation, as illustrated by recent Pacific Biosciences (PacBio) sequencing of 10×Genomics single cell libraries. While the accuracy of PacBio circular consensus reads facilitates cell barcode (cellBC) and unique molecular identifier (UMI) identification and thus assignment of a read to the correct cell and RNA molecule, the low PacBio throughput results in an insufficient coverage of the transcriptome (e.g. 270 reads, 260 UMI, 129 genes per cell in ^3^). This limits analysis to highly expressed transcripts. Oxford Nanopore Promethion long read sequencers generate at least 20 times more reads per flow cell than the PacBio Sequel II. However, the higher error rate of Nanopore sequencers renders cellBC and UMI identification challenging and has precluded so far the dissemination of Nanopore sequencing in high throughput scRNA-seq.

We have designed a strategy that enables efficient and reliable cellBC and UMI assignment to Nanopore reads. First, Illumina short read sequencing of the 10×Genomics libraries defines for each gene and genomic region the associated cellBCs and then for each cell/gene (or genomic region) the combination of associated UMIs. We subsequently used this information to guide cellBC and UMI assignment to the genome aligned Nanopore reads (Fig. 1a; detailed in Supplementary Fig. 1 and methods section).

**Figure 1.**
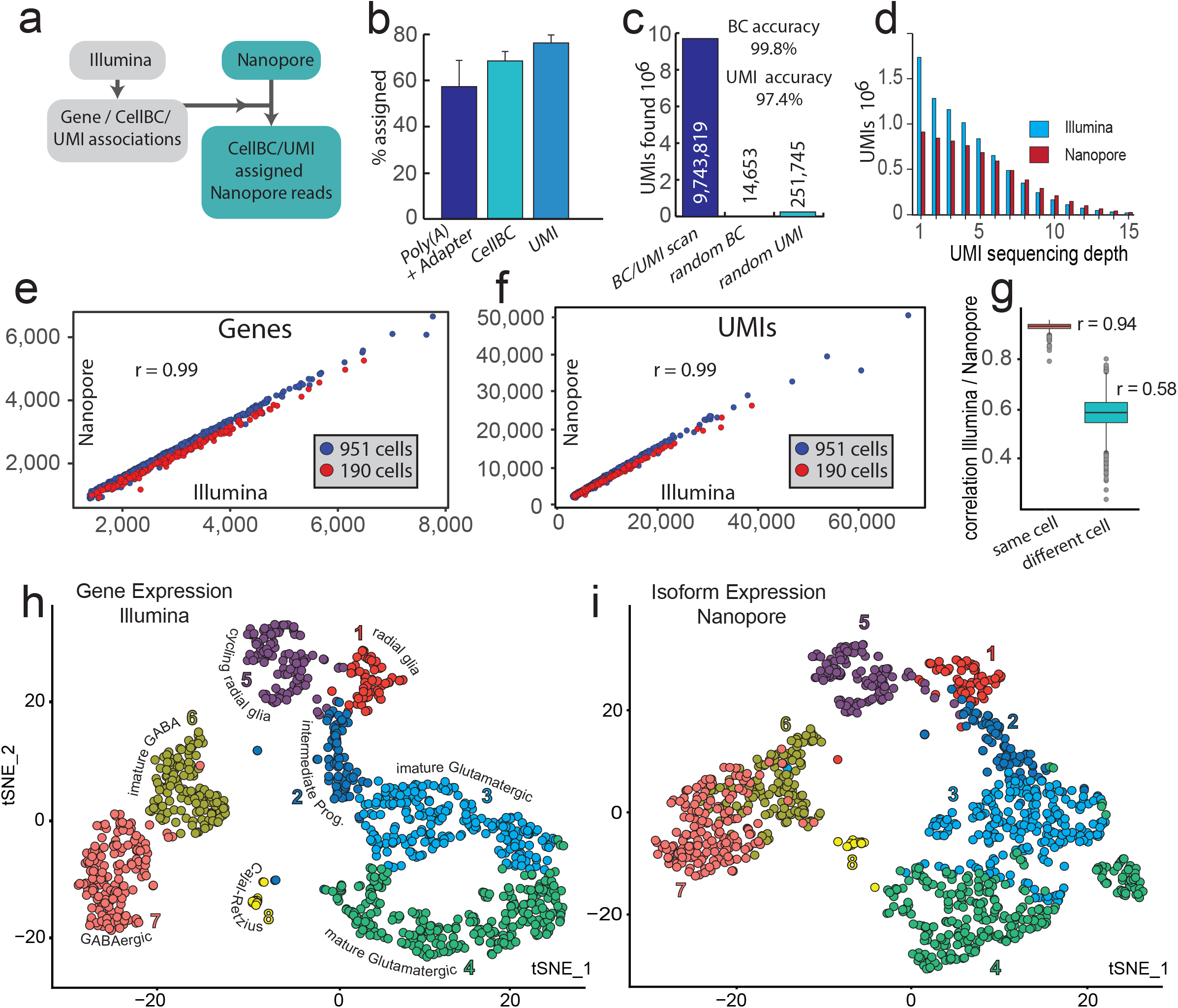
Nanopore scRNAseq. **(a)** Cell barcode and UMI assignment strategy (Detailed in Supplementary Fig. 1). **(b,c)** Efficiency (b) and accuracy (c) of cellBC and UMI assignment. **(b)** ly(A) and adapter, 57±11%; cellBC, 68±4%; UMI, 76±3%. **(c)** Number of UMI assigned reads before and after replacement of cellBC or UMI against random sequences (24×10^6^ reads scanned). **(d)** UMI sequencing depth (reads/UMI) for the Illumina and Nanopore dataset. **(e,f)**Number of genes (e) and number of UMIs (f) detected for each of the 1,141 cells after Illumina and Nanopore sequencing. **(g)** Correlation of gene expression between Illumina and Nanopore sequencing data for each cell. Boxes represent the 25% quantile to 75% quantile range. **(h,i)** t-SNE plots of Illumina gene expression (h) and Nanopore transcript isoform expression (i) for 1,141 cells. Colors for individual cells are the same in (h) and (i). Clustering details are in Supplementary Fig. 4 and Supplementary Table 1.

We prepared a 190 cell and a 951 cell E18 mouse brain library with the 10×Genomics Chromium system and generated 43 ×10^6^ and 70 ×10^6^ Illumina reads as well as 32×10^6^ and 322×10^6^ Nanopore reads for 190 and 951 cells, respectively. We identified poly(A) tails in 57 ± 11 % of the reads and assigned cellBCs with 99.8% accuracy to 67 ± 4 % of Nanopore reads with identified poly(A) sequence (Fig. 1 b,c; Supplementary Fig. 2). This scan eliminates most very low-quality reads since both poly(A) finding and barcode assignment are inefficient for low reads qualities (Supplementary Fig. 2). The poly(A) and cellBC discovery rates, as well as the accuracy of our software are, despite the higher error rates of Nanopore reads, similar to those reported previously for PacBio sequencing of 10×Genomics libraries^3^ and about tenfold higher than previously reported for Nanopore sequencing (identified cellBC in 6% of reads with poly(A) ^3^).

Molecular barcoding with UMIs is crucial for long read scRNA-seq since UMIs allow grouping of reads corresponding to the same RNA molecule, elimination of PCR amplification artifacts and correction of sequencing errors (Supplementary Note 1, Supplementary Fig. 3). In consequence UMIs minimize the risk that PCR generated chimeric cDNAs are falsely annotated as novel transcripts. However, previous studies failed to identify or did not use UMIs in single cell Nanopore sequencing ^3–5^. Our UMI assignment strategy in which we test for matching Illumina UMIs only for the same cell and the same gene or genomic region (detailed in Supplementary Fig1 and Methods section) strongly reduced the complexity of the UMI search and allowed, despite the high error rate of Nanopore reads, to assign UMIs with 97.4% accuracy to 76 ± 3 % of the reads with identified cellBC (Fig. 1b,c).

On average, 75% of the RNA molecules (UMIs) and 82% of the genes identified in each single cell after Illumina sequencing were also found in our Nanopore dataset (Fig. 1 e,f). The cellBC and UMI assigned Nanopore reads (median 28,120 /cell) reflect a median of 2,188 genes and 5,752 UMIs per cell with a good correlation between Nanopore and Illumina gene counts (Fig. 1e; r = 0.99), UMI counts (Fig.1f; r = 0.99) and gene expression for individual cells (Fig.1g, mean r = 0.94). In consequence, our cellBC and UMI assigned Nanopore dataset represents well the transcriptome captured in the 10×Genomics workflow. In total we found 18,439 Gencode v18 annotated and 15,317 novel transcripts backed by at least two UMIs that were each supported by at least 5 reads (Supplementary Note 2).

A t-SNE plot of the Illumina short read gene expression data reveals the cell types typical for E18 mouse brain (Fig. 1h). t-SNE projection of transcript isoform expression defined by Nanopore sequencing (Fig. 1i) yielded a similar clustering without revealing novel well defined sub-clusters. We next searched for genes with differential transcript isoform expression between clusters and identified cell type selective isoform usage for 139 genes and 324 differentially expressed isoforms (Supplementary Table 1, Supplementary Fig. 4b). For instance, Clathrin light chain A (Clta) (Fig. 2 a-d) and Myosin Light Chain 6 (Myl6; Supplementary Fig. 5) undergo a pronounced isoform switch during neuronal maturation. Myosin and Clathrin are involved in neuronal migration and axonal guidance in immature neurons^6^, ^7^ and in synaptic membrane recycling (Clathrin^8^) and synaptic remodeling (Myosin^9^) associated with synaptic plasticity in mature neurons. The isoform switch of Clta and Myl6 might fine-tune both proteins for their respective roles in different developmental stages.

**Figure 2.**
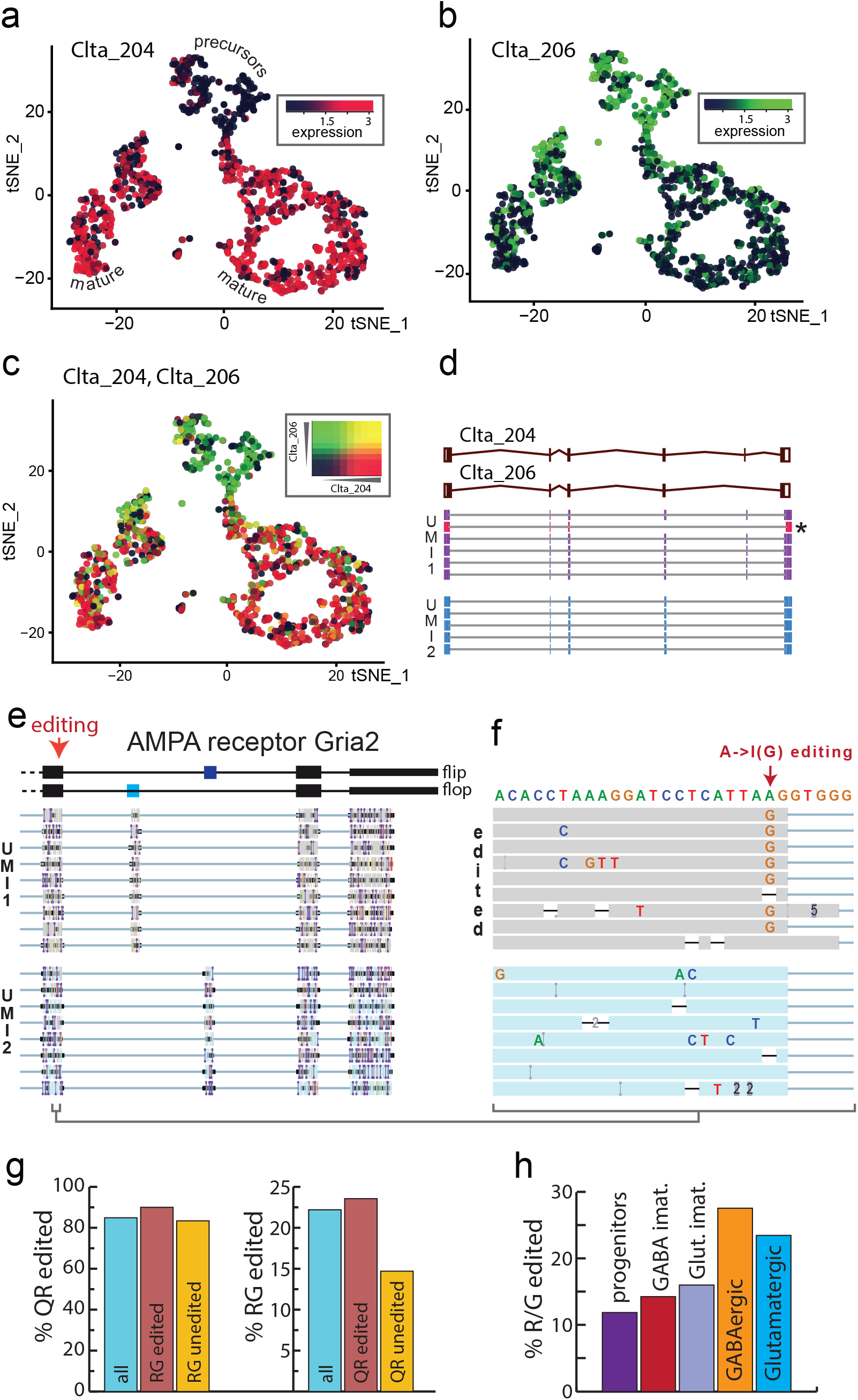
Nanopore scRNAseq reveals transcript diversity. **(a-c)** Clta isoform expression switch during neuronal maturation visualized on the t-SNE plot of Fig. 1e. **(a)** Clta-204 (ENSMUST00000107849.9), **(b)** Clta-206 (ENSMUST00000170241.7), **(c)** Overlay Clta-204 / Clta-206. **(d)** Principal Clta splice variants and supporting genome-aligned reads for two UMIs. A read not consistent with the UMI consensus (likely PCR artefact) is labelled with an “*”. **(e)** Principal splice variants of Gria2. The 3’ end of genome aligned reads for two UMIs are shown (Integrated Genome Viewer, UCSC Santa Cruz). **(f)** Zoom onto the R/G A->I editing site for the reads shown in (e). **(g)** Q/R and R/G editing. Editing rates at the Q/R site are 90% and 83.4% for RNAs edited or unedited at the R/G site respectively. R/G site editing is 23.6 % and 14.7 % for Q/R site edited and unedited RNAs respectively. **(h)** Extend of A->I editing of the R/G site increases during neuronal maturation. Progenitors, 11.9 %, clusters 1,2,5 in Fig. 1; GABA immature, 14.3 %, cluster 6; Glutamatergic immature, 16.0 %, cluster 3; GABA mature, 27.6 %, cluster 7; Glutamatergic mature, 23.5 %, cluster 4.

We next examined how single cell Nanopore sequencing with UMIs performs in defining SNVs for the ionotropic glutamate receptor Gria2, a post-synaptic cation channel that is A->I edited at two sites, leading to a Q/R substitution within the pore that renders the channel Ca^2+^ impermeable and a R/G substitution within the ligand binding domain that results in accelerated recovery from inactivation ^10^. Grouping of reads that corresponded to the same RNA molecule (UMI) allowed correction of Nanopore sequencing errors, generation of consensus sequence for each RNA molecule and identification of such sequence heterogeneity within single cells (Fig. 2 e,f). Analysis of error corrected Gria2 consensus sequences confirmed previous findings that Gria2 editing (Fig. 2g) is almost complete at the Q/R editing site (85 %) and partial at the R/G editing site (22 %). Long read sequencing further revealed that editing of one site increases the probability that the other site is edited (Fig. 2g) and that editing of the R/G site increases during neuronal maturation (Fig. 2h) from 11 % in neuronal progenitors to 27.5 and 23.5% in mature inhibitory and glutamatergic neurons respectively. Thus, single cell long read sequencing both confirms and extends previous knowledge on Gria2 editing in the central nervous system.

In conclusion, combining the high throughput of Nanopore sequencing with UMI guided error correction allows both high confidence definition of transcript isoforms and identification of sequence heterogeneity in single cells. Single cell Nanopore sequencing with UMIs (ScNaUmi-seq) will facilitate high throughput single cell studies on RNA splicing, editing and imprinting and should also become a valuable tool for in depth studies of tumor heterogeneity.

## METHODS

### Mouse brain dissociation

A combined hippocampus, cortex, and ventricular zone pair from an E18 C57BL/6 mouse was obtained from BrainBits LLC (Leicestershire, UK). A single cell suspension was prepared following the 10×Genomics protocol for “Dissociation of Mouse Embryonic Neural Tissue”. Briefly, tissues were incubated with 2mg/ml Papain in calcium free Hibernate E medium (BrainBits) for 20 min at 37°C, rinsed with Hibernate-E/B27/GlutaMAX (HEB) medium, triturated and filtered through a 40 μm FlowMi cell strainer (Sigma-Aldrich). Cell concentration and viability was determined with a Countess® II automated cell counter (Life Technologies).

### Single cell cDNA library preparation

The E18 mouse brain single-cell suspension was converted into a barcoded scRNA-seq library with the 10×Genomics Chromium Single Cell 3’ Library, Gel Bead & Multiplex Kit and Chip Kit (v2), aiming for 1.400 cells and following the manufacturer’s instructions with the following modifications: To obtain batches with different cell numbers, we split the emulsion before reverse transcription into two aliquots (85.7% and 14.3%; targeting 1200 and 200 cells). We extended the PCR elongation time during the initial PCR amplification of the cDNA from the manufacturer recommended 1 min to 3 min to minimize preferential amplification of small cDNAs. Half of the amplified cDNA was used for short read sequencing library preparation following the 10×Genomics protocol and sequenced on an Illumina Nextseq 500 sequencer (26bases + 57bases). We generated 43M and 70M Illumina reads for the samples targeting 200 and 1200 cells respectively and mapped them to the mouse genome (build mm10) with the 10×Genomics CellRanger software (v2.0.0).

For each Promethion flow cell (Oxford Nanopore) we re-amplified 2 - 10 ng of the 10×Genomics PCR product for 8 cycles with the primers NNNAAGCAGTGGTATCAACGCAGAGTACAT and NNNCTACACGACGCTCTTCCGATCT (Integrated DNA Technologies, IDT). The Ns at the 5’ end of the primers avoid the preferential generation of reverse Nanopore reads (> 85%) we observed without those random nucleotides. Amplified cDNA was purified with 0.65x SPRISelect (alternatively: 0.45× Spriselect for depletion of cDNA smaller than 1 kb) and Nanopore sequencing libraries were prepared with the Oxford Nanopore LSK-109 kit following the manufacturer’s instructions. For the libraries targeting 200 and 1200 cells we generated 32M reads and 322M reads respectively.

#### Optional steps for the depletion of cDNA lacking a terminal poly(A) / poly(T)

Amplified single cell cDNA contains to a variable extend (30 – 50 %) cDNA that lacks poly(A) and poly(T) sequences. For the study presented here, we did not deplete those cDNAs. Such cDNA can be depleted after a PCR of 2 – 10 ng of the 10×Genomics PCR product for 5 cycles with 5’- NNNAAGCAGTGGTATCAACGCAGAGTACAT-3’ and 5’ Biotine-AAAAACTACACGACGCTCTTCCGATCT 3’. After 0.55× SPRIselect purification to remove excess biotinylated primers, biotinylated cDNA (in 40 μl EB) is bound to 15 μl 1× SSPE washed Dynabeads™ M-270 Streptavidin beads (Thermo) resuspended in 10 μl 5× SSPE for 15 min at room temperature on a shaker. After two washes with 100 μl 1× SSPE and one wash with 100 μl EB, the beads are suspended in 100 μl 1× PCR mix and amplified for 6 – 9 cycles with the primers NNNAAGCAGTGGTATCAACGCAGAGTACAT and NNNCTACACGACGCTCTTCCGATCT to generate enough material for Nanopore sequencing library preparation.

All PCR amplifications for Nanopore library preparations were done with Kapa Hifi Hotstart polymerase (Roche Sequencing Solutions): initial denaturation, 3 min at 95°C; cycles: 98°C for 30 sec, 64°C for 30 sec, 72°C for 5 min; final elongation: 72°C for 10 min, primer concentration was 1 μM.

### Mapping of Nanopore reads

Nanopore reads were aligned to the *Mus musculus* Genome (mm10) with minimap2 v2.17 in spliced alignment mode (command: ‘minimap2 -ax splice -uf --MD -N 100 --sam-hit-only --junc-bed’. The splice junction bed file was generated from the Gencode v18 GTF using paftools.js, a companion script of minimap2. For reads matching known genes, the gene name was added to the corresponding SAM record (SAM Tag: GE) using the Drop-seq tools package (v1.12) ^11^. Before cellBC and UMI assignment, SAM records were annotated with their Nanopore read sequence (SAM Tags: US) and read qualities (SAM tag: UQ).

### CellBC and UMI assignment to Nanopore reads

Our Java software performs the following analysis steps (Suppl. Fig. 1).

#### Parsing of Illumina data

To retrieve accurate cellBC and UMI information, BAM files with the cellBC and UMI assigned Illumina data generated by the 10×Genomics Cellranger software were parsed. For each gene, we extracted the cellBC from the Illumina data and identified the UMIs found for each gene/cellBC combination. We also associated genomic regions (window size 500 nt.) with cellBCs and UMIs to account for reads that match outside of annotated genes. The parsed Illumina data were stored in nested Hashtables as serialized Java objects.

#### Search for poly(A) tail

Our software searches for a poly(T) and a poly(A) sequence (default 85% A or T, ≥ 20 nt.) within 100 nucleotides from the 5’ or 3’ end of the read, respectively. Reads without poly (A or T) and reads with a poly(A or T) on both ends are not further analyzed.

#### Search for 3’ adapter sequence

The cellBC and the UMI are located between an adapter sequence (10×Genomics 3’ PCR priming site) and the poly(T) of the reverse transcription primer (Supplementary Fig. 1 b). To define the position of the cellBC we searched for the adapter sequence between the extremity of the read and the poly(A/T) sequence identified in the previous step using sliding window Needleman Wunsch alignments. The position with the best adapter match (least mismatches) was used. We found that searching for just the ten 3’ nucleotides of the adapter with 3 allowed mismatches (adapter found in 90.4 % of the reads with poly(A)) more efficient than a search for a 20 nucleotide adapter sequence with 6 allowed mismatches (79.9% adapter found in poly(A) reads). Possible reasons for this are (i) The adapter 5’ end is very close to the extremity of the read and read quality might be lower there. (ii) Intrinsic error rate of Nanopore sequencing might be higher for the 5’ of the adapter sequence.

#### Search for internal adapter and poly(A)

To flag reads corresponding to chimeric cDNA generated during library preparation, we searched for internal adapter sequences in proximity of a A- or T-rich sequence and flagged those reads as chimeric reads in the output file. In our dataset we found internal adapters in 3.5% of the reads.

#### Search for cellBCs

Cell barcodes in high accuracy Illumina reads are typically assigned by grouping reads that differ by not more than one position (edit distance: ED = 1). Indels are typically not considered. In consequence just 48 possible permutations of the 16 nt. cellBC reads need to be analyzed and assignment of reads to a barcode is highly reliable. Nanopore reads still have a mean error rate of about 5 – 10 % (substitutions and indels). In consequence higher edit distances need to be examined and indels must be considered. This implies the generation and analysis of 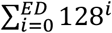 barcode sequence permutations (2,113,664 for ED=3) for each read. A 50 million read PromethION sequencing run would require the generation and analysis of about 10^14^ barcode sequences for ED=3. This is clearly not feasible using reasonable sized compute clusters and standard bioinformatics approaches where sequences are typically treated as text.

To solve this computational bottleneck we encoded the barcode sequences using 2 bits per base (A: 00, G:01,T:10,C:11) which allows encoding of the entire 16 nucleotide cellBC into just one integer. This bitwise encoding allows performing substitutions, insertions and deletions using highly efficient bitwise operations that require just one CPU cycle. Encoding cellBC and UMIs into integers also tremendously accelerates the search for matching Illumina barcodes or UMIs, since searching for a matching integer is much faster than searching for matching strings.

An additional challenge for accurate barcode assignment is that 10×Genomics cellBCs are randomly selected out of a pool of 750,000 barcode sequences. The used barcode sequences are not known in advance. Clustering the Nanopore barcode reads correctly without a priori knowledge of the used barcodes is rather error prone, since two reads that have e.g. two sequencing errors in the cellBC each, can differ in up to four positions when compared to each other.

To improve the accuracy of barcode assignment, we used the cellBC sequences defined by Illumina sequencing of the same libraries to guide the cellBC identification in Nanopore reads. For each genome aligned Nanopore read, we extracted the sequence (16 nt. barcode plus 4 nt. to allow insertion errors) just downstream of the adapter (position corrected for indels in adapter) identified in the previous step. We also extracted the barcode sequences for the preceding and following position to account for eventual terminal indels in the adapter read. We then compared the extracted Nanopore barcode sequences and all possible permutation up to a defined edit distance with: (i) The cellBCs identified by Illumina sequencing for the same gene if the Nanopore read matches a known gene in the Illumina dataset; (ii) Illumina cellBCs sequences for the genomic region (500 nt. window) if the Nanopore read matches an unannotated genomic region or if the gene name was not found in the Illumina data. Since the Nanopore barcodes are only compared with Illumina barcodes found for the same gene or genomic region, the complexity of the search set is reduced. Barcode matches, eventual second-best barcode matches and their quality (edit distance) were recorded in the output BAM file. The maximal ED is dynamically selected (ED limit 1-4) and depends on the number of cellBCs found for the same gene or genomic region in the Illumina data (the complexity of the search set). The software allows to define the maximal false assignment percentage as a parameter. We used computer simulations of collision frequencies to define the complexity of the search set (number of Illumina barcodes) allowed for different edit distances and maximal false assignment percentages. Simulation data are supplied as an XML file and can be easily adapted. Optionally the software also allows the definition of fixed edit distance limits.

#### Search for UMIs

To identify the ten nucleotide UMI, 14 nucleotides (to allow for insertions in the read) following the end of the cellBC sequence (position corrected for barcode indels) were extracted. For UMI assignment, we used the same strategy as for cellBC assignment. We searched for matching UMIs in Illumina data for the same cellBC and genomic region or gene identified in the previous step. This means the complexity of the search set corresponds to the copy number of a given transcript in just one cell. The maximal edit distance allowed is dynamically adjusted depending on the complexity of the search set.

#### Determination of cellBC and UMI assignment accuracy

To evaluate the accuracy of cellBC and UMI assignment, we first scanned each Nanopore SAM record for a matching Illumina cellBC or UMI. We then repeated the scan where we replaced each cellBC or UMI sequence extracted from the Nanopore read by a random sequence and mutated this random sequence allowing the same number of mutations (edit distance) that was used for the same SAM record in the previous scan and searched for matching cellBC or UMIs in the Illumina dataset. The ratio between the number of reads that were assigned to both a cellBC and UMI after and before replacement of the cellBC or the UMI against a random sequence is the false assignment probability.

The cellBC assignment accuracy is particularly high, since after an incorrect cellBC assignment, the UMI is compared with the UMIs associated with this wrong cellBC in the Illumina data. In consequence, most reads with falsely assigned cellBC are subsequently eliminated during UMI assignment. The false barcode assignment rate in cellBC and UMI assigned reads is thus the product of the false cellBC and the false UMI discovery rate. With the default cellBC (5% false assignment) and UMI scanning (2% false assignment) parameters we used; we obtained an effective cellBC assignment accuracy of 99.8% what is close to the value expected with those parameters.

#### Maximal possible Barcode and UMI assignment efficiency with 10×Genomics data

Since our software scans for matches of the Nanopore cell barcodes with barcodes associated with cells (the relevant barcodes), barcodes associated with empty drops are ignored. The maximal possible barcode discovery rate in Nanopore reads corresponds to the percentage of reads associated with cells in the Illumina dataset: 83.7% and 85.1% for the 200 cell and the 1200 cell sample respectively.

The maximal UMI discovery rate for Nanopore reads depends on the Illumina sequencing depth of the same sample and corresponds to the sequencing saturation computed by the 10×Genomics CellRanger software after Illumina sequencing. The sequencing saturation is the probability that a matching UMI is found in the Illumina dataset for a given read. For the 190 cell and 951 cell samples, the sequencing saturations were 90.5% and 74.8% respectively. Efficiencies of barcode assignment are given as percentages of cell associated barcodes. UMI assignment efficiencies were corrected for the sequencing saturation.

#### Compatibility of the software

The software is compatible with the 10×Genomics workflow v2 and the recent upgrade (v3) which uses 12nt UMIs. It can also be used for cellBC and UMI assignment of long read single cell data generated with other single cell isolation systems with the following limitations: The cDNA needs to have a 3’ adapter followed by a cellBC, an UMI and a poly(A). CellBC and UMI length, adapter sequences as well as the search stringency for poly(A/T), adapter, cellBC and UMI can be configured accordingly.

### Assignment of UMIs to known transcripts

SAM records matching known genes were grouped by UMI and analyzed for matching Gencode v18 transcripts. SAM records with extensive non-matching sequences at either end (hard- or soft clipping > 150 nt.) were discarded (1,93 % and 3,95% for the 190 and 951 cell dataset respectively). To assign a UMI to a Gencode transcript we used the following strategy: For every UMI each genome mapped read was examined for exon junctions matching the junctions present in the Gencode transcripts for the corresponding gene, authorizing a two bases margin of added or lacking sequences at exon boundaries, to allow for indels at exon junctions and imprecise mapping by Minimap2. For each found exon-exon junction, a score of 1 was added to each Gencode isoform carrying the junction. Finally, if just one isoform had the highest score it was selected as the isoform for the UMI. Following this strategy, we assigned 77.9 % of the UMIs to a known Gencode transcript isoform.

### Quantification of novel transcript isoforms

Consensus sequences for UMIs (1,141 cells dataset) backed by at least 5 nanopore reads were used for quantification of novel transcripts with SQANTI2 (https://github.com/Magdoll/SQANTI2). Consensus cDNA sequences for UMIs backed by at least five reads were mapped to the mouse genome (mm10) with minimap2 (« -ax splice -t 6 -uf --secondary=no -C5»). UMIs were then grouped by transcript isoform (exon makeup) with the “collapse_isoforms_by_sam.py” script (https://github.com/Magdoll/cDNA_Cupcake, parameters: « --dun-merge-5-shorter --max_fuzzy_junction 0 -c 0.95»). Isoforms supported by less than five UMIs were discarded. Isoforms were then compared to Mouse Gencode v18 and classified and counted with Sqanti2^12^ using default parameters. The number of known Gencode transcripts found in our dataset are the transcripts classified by Sqanti2 as “Full Splice Match” (all and only exon-exon junctions of the reference are present). Novel transcript isoforms are transcripts with an exon makeup not consistent with the Gencode reference. To define the number of novel transcripts we used the union of transcripts classified by Sqanti2 as “Novel In Catalogue”^12^ and the Sqanti2 category “Novel Not In Catalogue”^12^. We next removed mono-exonic transcripts and filtered for transcripts with all exon-exon junctions either described in Gencode or confirmed in E18 cortex and midhindbrain Illumina short read data (Geo accession GSE69711).

### Single-cell gene expression quantification and determination of major cell types

Raw gene expression matrices generated by CellRanger were processed using R/Bioconductor (version 3.4.3) and the Seurat R package (version 3.0.0). A total of 190 cells and 951 cells were detected with default CellRanger cutoffs for the two datasets respectively. Cells with over 95% dropouts were removed. From the 186 and 935 remaining cells, gene expression matrices were cell-level scaled to 10.000 and log-normalized. The top 2,000 highly variable genes were selected based on the variance-stabilizing transformation method and used for the Principal Component Analysis. Due to differences in sequencing depth of both datasets, data were integrated using the Seurat CCA method. The first 11 aligned canonical correlations were used to define the integrated sub space for clustering and tSNE visualization of the 1,121 remaining cells. Clusters in the tSNE plot were assigned to known cell types using canonical marker genes (Supplementary Fig. 4a, Supplementary Table 1).

### Analysis of Nanopore single cell gene and isoform expression data

For cellBC/UMI assigned Nanopore data, we generated isoform-level (median UMIs/cell = 4,472) and gene-level (median UMIs/cell = 5,752) count matrices using custom software (IsoformMatrix function of Sicelore-1.0.jar) and processed the data with the Seurat R package (version 3.0.0). Nanopore gene expression data were used for the Nanopore/Illumina gene count and UMI count correlations in Fig. 1. Nanopore isoform data were subjected to the same statistical analysis as the Illumina short reads to generate the Isoform-level tSNE representation (Fig.1b).

### Consensus sequence generation

For each UMI, we aligned the corresponding reads and called the consensus sequence with poaV2 ^13^. The consensus sequences were polished with racon ^10^ using all reads available for the UMI. Consensus sequences were then mapped again to the genome using Minimap2 and the resulting BAM file was subsequently used for SNP calling (e.g. analysis of RNA editing). When just two reads were available for a UMI, the read with the best reference genome match was used.

### Gria2 data analysis

9,593 reads (2,105 UMIs) corresponding to Gria2 (mm10:chr3:80,682,936-80,804,791) were extracted from the 951 and the 190 cell dataset. A consensus sequence for each molecule was computed and re-mapped to the mm10 genome for SNP calling. Gria2 mRNAs are huge (> 6 kB) and inefficiently converted into full length cDNA in the 10×Genomics workflow. This is likely due to: (i) some RNA degradation within the droplet between cell lysis and reverse transcription. (ii) internal reverse transcription priming at A-rich sites within the cDNA leading to cDNAs that cover only part of the transcripts. In consequence, we noticed 3’ bias and fragmented coverage for certain long transcripts such as Gria2 where a total of 456 cDNA molecules (UMIs) had the R/G-editing-and 225 had both the R/G and the Q/R-editing-site. Further optimization of the 10×Genomics workflow will be required for more efficient full-length capture of long mRNAs.

## DATA AVAILABILITY

All relevant data have been deposited in Gene Expression Omnibus under accession number GSE130708 (https://www.ncbi.nlm.nih.gov/geo/query/acc.cgi?acc=GSE130708, token: arwjecaifvcxrsh).

## CODE AVAILABILITY

All custom software used is available on Github https://github.com/ucagenomix/sicelore.

## ACKNOWLEDGEMENTS

This project was funded by grants from the Conseil Départemental des Alpes Maritimes (2016-294DGADSH-CV), FRM (DEQ20180339158), the Chan Zuckerberg Initiative (Silicon Valley Foundation, 2017-175159-5022), the National Infrastructure France Génomique (Commissariat aux Grands Investissements, ANR-10-INBS-09-03, ANR-10-INBS-09-02).

## SUPPLEMENTARY INFORMATION

### SUPPLEMENTARY FIGURES

**Supplementary Figure 1.**
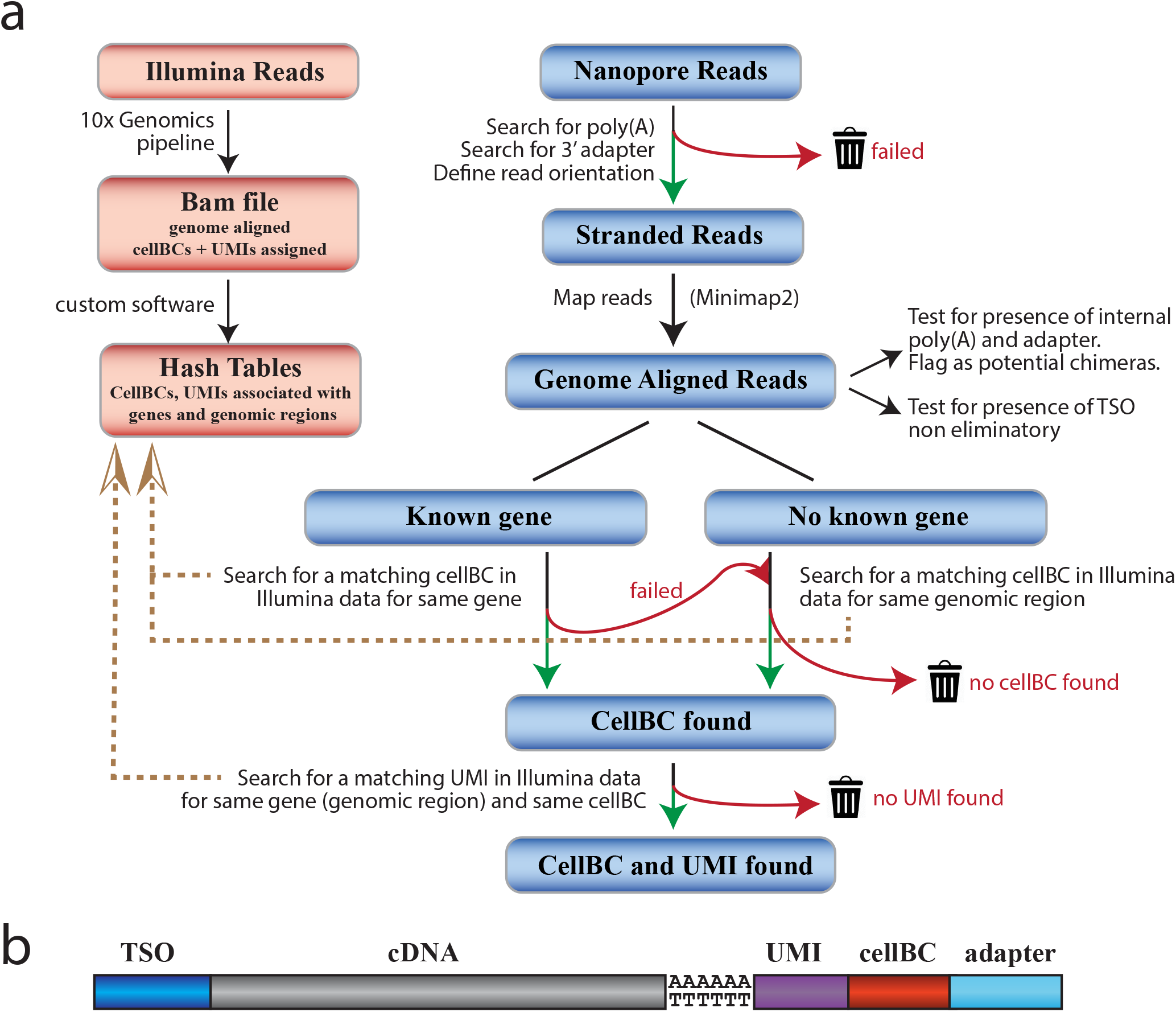
CellBC and UMI assignment strategy. **(a)** CellBC and UMI assignment strategy. See methods section for description of the individual steps. **(b)** Organization of single cell cDNA generated by the 10×Genomics workflow.

**Supplementary Figure 2.**
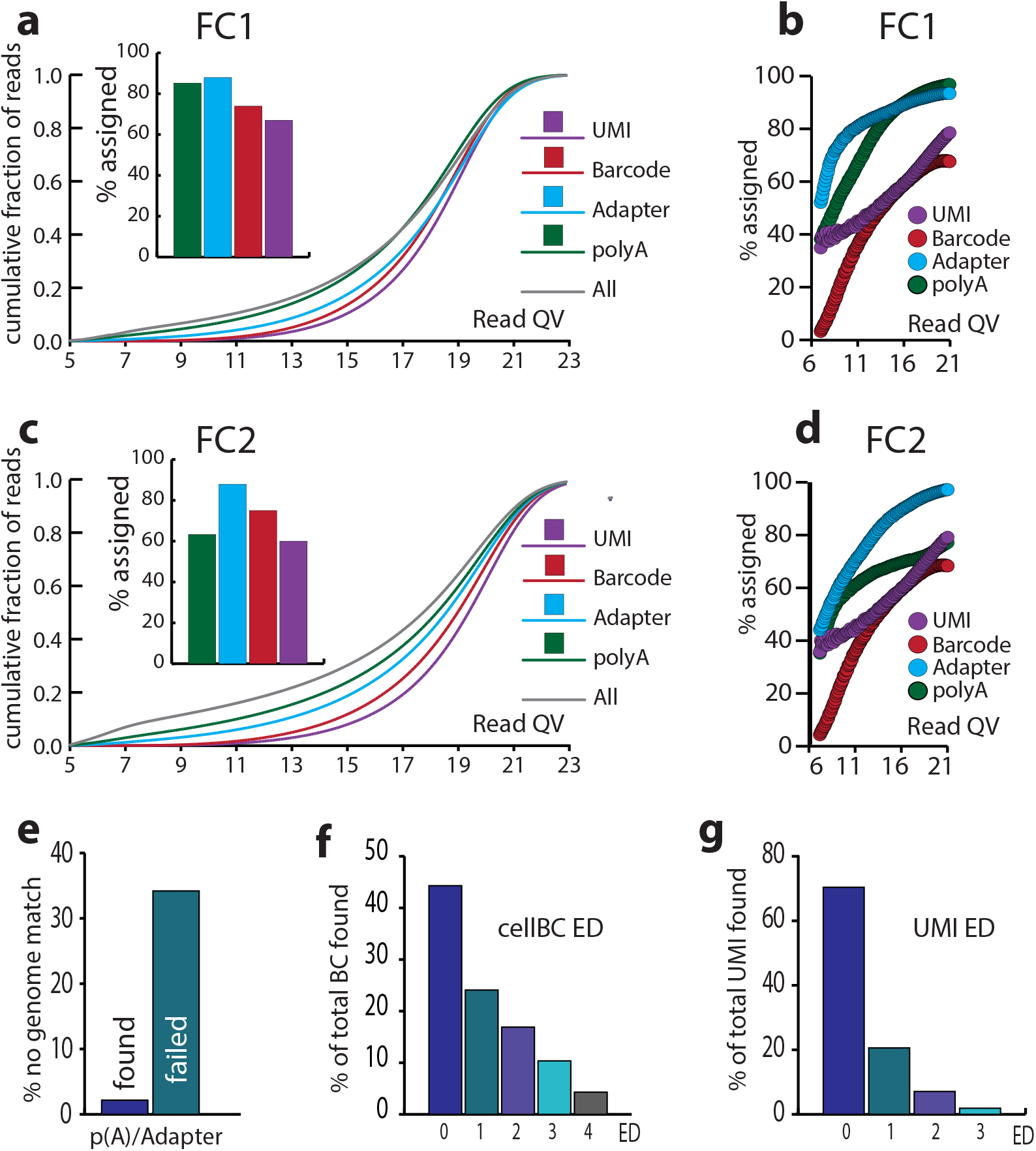
Efficiency of cellBC and UMI assignment. (**a,c**) Cummulative read quality distributions for two representative Promethion sequencing runs (a: FC1, 32 *10^6^ reads, 190 cells; c: FC2 70 *10^6^ reads, 951 cells) are shown for all reads (grey), polyA- (green), adapter- (light blue), cellBC- (red) and UMI- (purple) assigned reads. Fractions of reads (%) in windows of 0.1 (PHRED score) are shown. Histograms show fractions of reads that passed the poly(A) scan, adapter scan, cellBC assignment, UMI assignment. **(b,d**) Relation between read quality and poly(A), adapter, cellBC and UMI assignment efficiency. Dashed vertical lines indicate mean QV of all reads. Poly(A), adapter, cellBC and UMI assignment efficiencies in (a-d) are relative to total reads, adapter assigned reads and cellBC assigned reads respectively. **(e)** The poly(A) scan eliminates the majority of reads that don’t match the genome. Fraction of unmapped reads with found poly(A) and adapter (2.2% of 24,008,457) and for reads without identified poly(A) and adapter (35.1% of 8,062,431) for FC1. **(f,g)** Edit distance distribution (mismatches between Nanopore and Illumina) for found cellBC (f) and UMIs (g). Shown are % of total found cellBCs or UMIs for each edit distance.

**Supplementary Figure 3.**
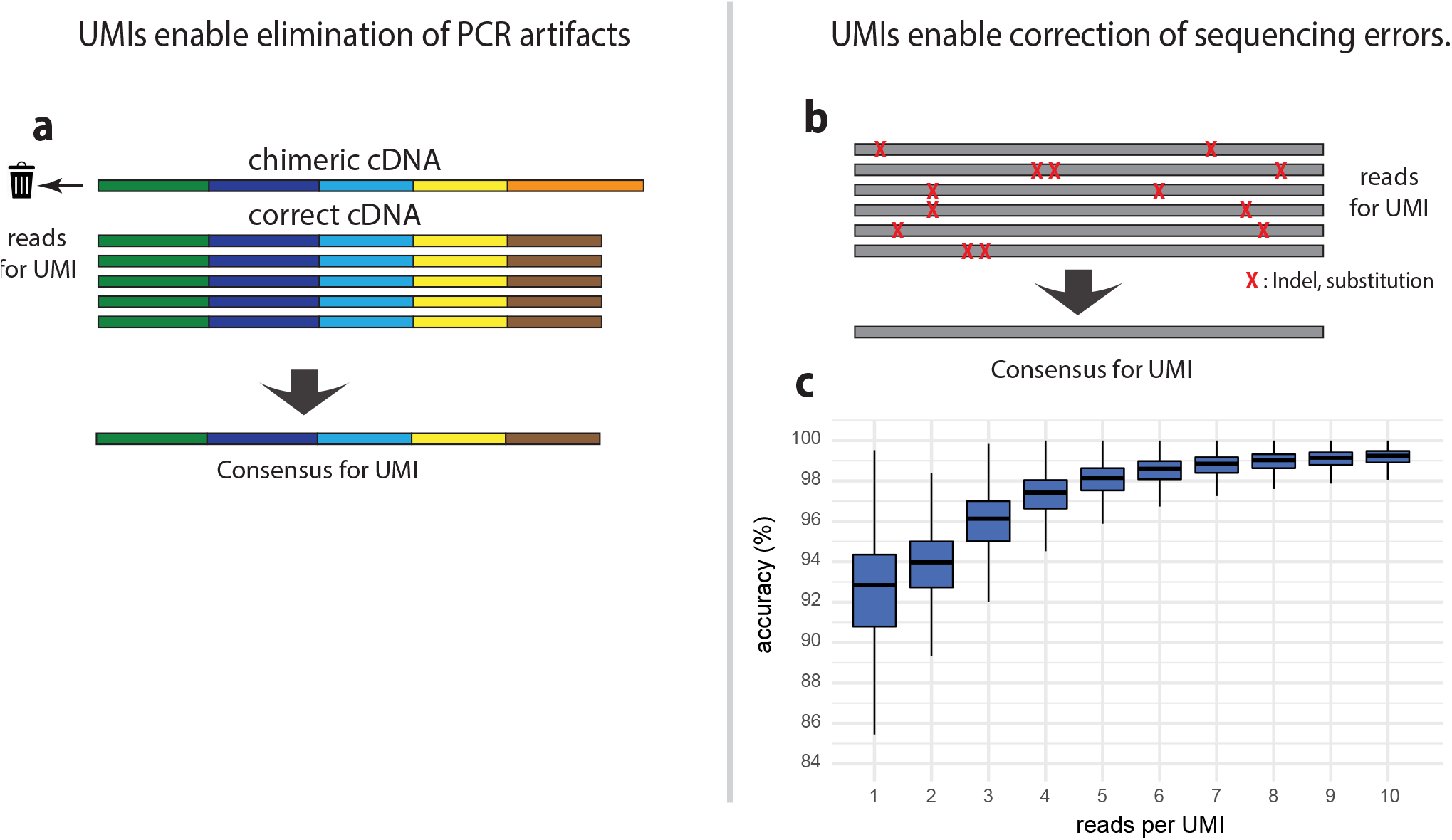
UMIs enable identification of PCR artefacts and sequencing error correction. **(a)** UMIs allow identification and elimination of reads originating from chimeric cDNA generated during PCR amplification. Chimeric cDNAs are mainly generated in later PCR cycles when cDNA concentration becomes higher. This results in a small fraction of reads (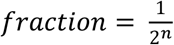, n : PCR cycle where chimera is generated) with aberrant exon layout. Those inconsistent reads are discarded, and the remaining reads are used to define the consensus cDNA sequence for the UMI. **(b, c)** UMIs enable sequencing error correction. Generation of consensus sequences for all reads associated with each UMI allows to obtain high accuracy sequences for each UMI (RNA molecule) despite the low accuracy of each Nanopore read. **(c)** the box plots indicate the accuracy of the cDNA consensus sequences (% identity to the reference genome) for different UMI sequencing depths. Boxes represent the 25% quantile to 75% quantile range, upper and lower edges of notches are median +/− 1.58 * IQR / sqrt(n) (IQR : inter quantile range, n: sample size).

**Supplementary Figure 4.**
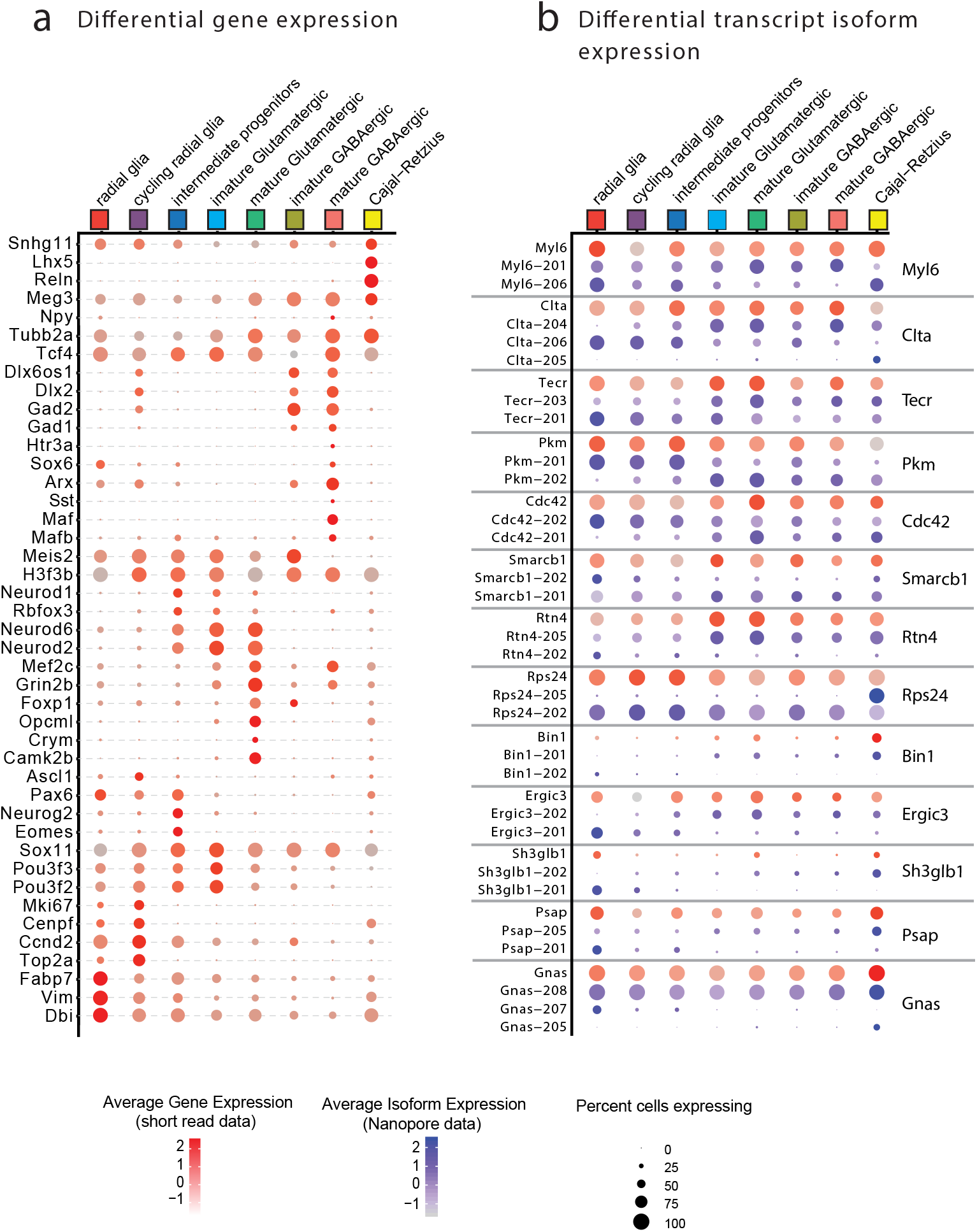
Differential gene and transcript isoform expression in subtypes of neurons. **(a,b)** Expression of selected genes (a) and isoforms (b) in cell types (Fig. 1e). **(a)** Illumina short read sequencing gene expression for a subset of the genes markers differentially expressed between clusters. **(b)** Gene (red dots) and isoform (blue dots) expression for selected genes that show differential expression of isoforms between neuronal subtypes (clusters). Dot size and color intensity represent the percentage of expressing cells in cluster and expression level (ln(count)) respectively. Transcript names are Ensembl names. The full list of differentially expressed genes and transcript is in Supplementary Table 1.

**Supplementary Figure 5.**
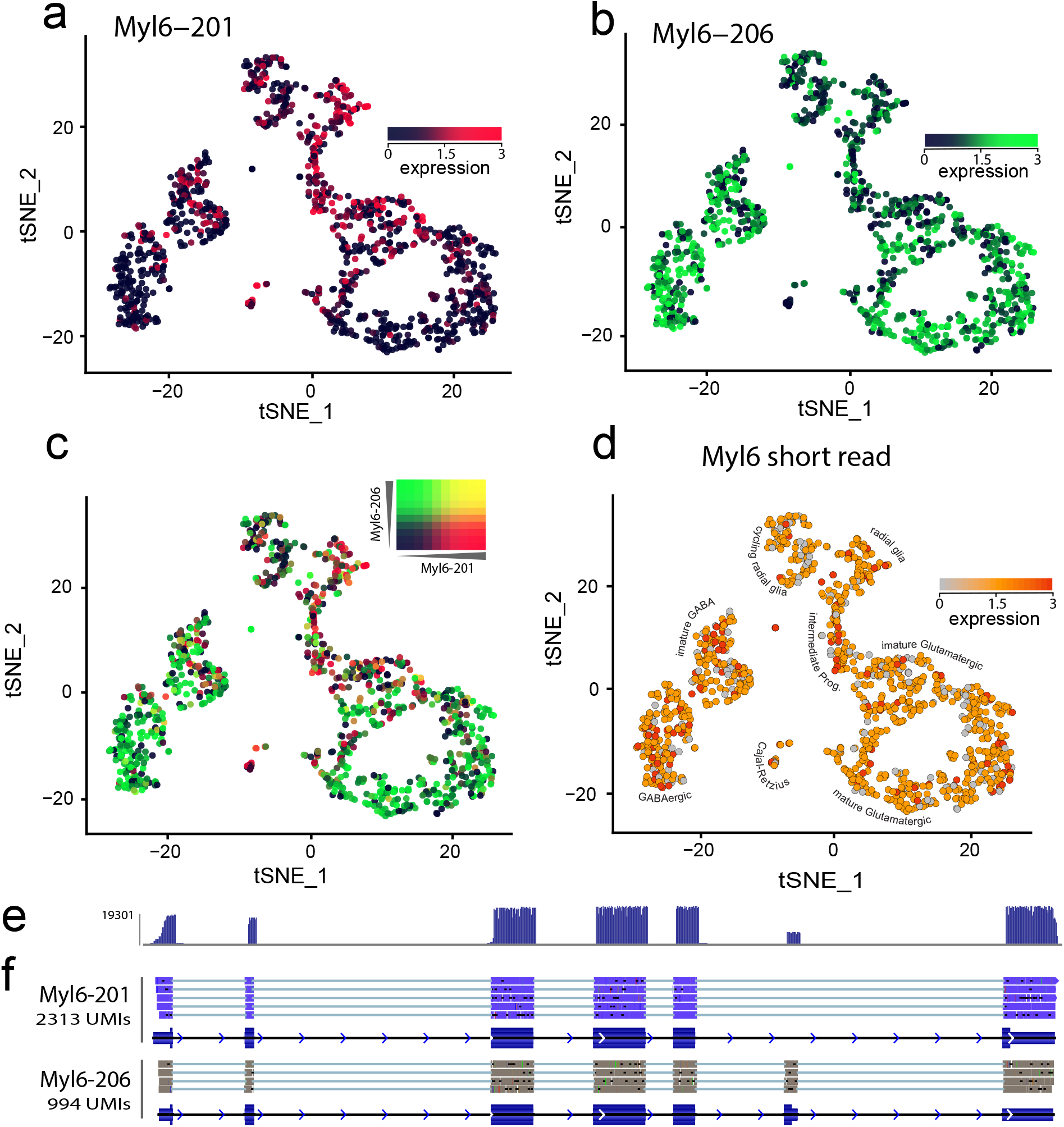
Myl6 alternate splice variants are differentially expressed during neuronal maturation. **(a-c)** Isoform switch of the essential (alkaline) Myosin light chain 6 during neuronal maturation visualized on the t-SNE plot of Fig. 1e (1141 cells). (a) Myl6-201 (ENSMUST00000164181.1), (b) Myl6-206 (ENSMUST00000218127.1), (c) Overlay Myl6-201/ Myl6-206. **(d)** Myl6 gene expression from Illumina short read data. **(e)** Intron/exon structure of the two principal Myl6 splice variants. Filled bars represent coding regions. **(f)** Nanopore read density distribution over the Myl6 gene. **(g)** Genome aligned Myl6 reads for two UMIs. The number of UMIs is indicated for each isoform. (f) and (g) are screenshots of the aligned Nanopore Bam files visualized with IGV (Integrated Genome Viewer, UCSC Santa Cruz).

### SUPPLEMENTARY NOTES

#### Supplementary Note 1: Unique molecular identifiers are crucial for accurate long read single cell sequencing

In single cell transcriptome sequencing a few picograms of mRNA in each single cell is converted into cDNA and heavily PCR amplified during sequencing library preparation. However, PCR amplification has several well-known issues such as the generation of chimeric cDNA that might get classified as novel transcript variants or fusion transcripts.

Tagging each cDNA with a “Unique Molecular Identifier” (UMI) during reverse transcription allows grouping of reads that originate from the same RNA molecule and to eliminate such PCR generated chimeric cDNAs. Reads from PCR artifacts typically represent a minority of the reads of a UMI and are not consistent with the other reads for the same UMI (Supplementary Fig. 3 a). Elimination of PCR artifacts with this strategy requires that UMIs are sequenced several times. In consequence, a high sequencing depth is crucial. Currently two high throughput long read systems are commercialized, the Pacific Biosciences Sequel II and the Oxford Nanopore Promethion. While PacBio circular consensus sequences (CCS) have higher accuracy (99%) than Nanopore reads (93% mean accuracy), the number of reads generated by a PacBio Sequel II flow cell (2 million CCS) is too low for a sufficiently high sequencing depth at a reasonable cost. The Oxford Nanopore Promethion generates in our hands > 70 million reads per flow cell. The principal weakness of Nanopore sequencing, the lower accuracy, can be addressed by UMIs. PacBio achieves a high sequencing accuracy despite the raw read accuracy of just 85% by performing multiple sequencing rounds on circularized library DNA and generation of a consensus sequence from those reads. Grouping of multiple reads for each UMI allows to achieve a similar accuracy boost for Nanopore reads and enables the generation of high accuracy consensus sequences for each RNA molecule (Supplementary Fig. 3 b,c).

The principle unsolved challenge that prevented the use of UMIs in Nanopore sequencing is the accurate identification of the 10 nt. UMI sequences in low accuracy Nanopore reads.

For PacBio scRNAseq, Gupta et al.^3^ used a strategy where they grouped identical UMI read sequences. The drawback of this strategy is that each UMI read that has an error yields a novel false UMI. With the 1% PacBio and the > 5% Nanopore sequencing error rate, 10% and 50% respectively of the 10 nt. UMI reads are expected to have at least one error and to yield a novel wrong UMI.

A more robust UMI assignment strategy that is less sensitive towards sequencing errors is required to enable the use of UMIs in single cell Nanopore sequencing.

We developed a UMI assignment strategy (Methods section and Supplementary Fig. 1) that tolerates sequencing errors where we compare the Nanopore UMI reads with the UMIs defined with high accuracy for the same gene and the same cell by Illumina sequencing (see Methods section). Since with this strategy, a UMI matching the Nanopore read is searched within just a small set of UMIs, we assign UMIs in Nanopore reads with high accuracy (97.4%, Fig. 1c).

#### Supplementary Note 2: Identification of novel transcript isoforms

Single cell long read sequencing is a powerful approach to detect cell type selective expression of transcript isoforms. However, before rare novel sequences are coined as novel transcript isoforms potential artifacts that are ixsnherent to most common RNA-seq library preparations that involve RT and PCR amplification need to be considered. Both RT and PCR generate artifactual cDNA. While PCR artifacts can be eliminated when UMIs are used (Supplementary Note 1), identification of RT artifacts is, in general, more difficult. Reverse transcriptases tend to switch templates between homologous sequences^1^ resulting in deletions (intramolecular) or chimeric cDNA (intermolecular).

When RNAseq libraries are prepared from purified RNA, the sample is denatured to remove RNA secondary structure, snap cooled, and RT is carried out at ≥ 37°C. In scRNAseq with the most widely used systems (e.g. 10× Genomics Chromium), cells are co-encapsulated with RT reagents and RT-primer beads into emulsion droplets where cells are lysed. Thus, reverse transcription is initiated at room temperature without disrupting the RNA secondary structure and proceeds at low stringency until the emulsion is recovered and heated to complete the RT. Low temperatures during RT, as well as inter and intramolecular secondary structure of RNA, is known to favor RTase template switching^14^. The low stringency initiation of the RT within the droplets also leads to false priming of the oligo-dT primers on A-rich sequences. We observe such RT priming on A-rich sequences mainly within 3’ UTRs or intronic sequences of partially spliced mRNAs. Thus, single cell cDNA has typically a higher fraction of artifactual cDNA than cDNA prepared from purified mRNA under well controlled conditions.

To evaluate the number of novel transcript isoforms in our scRNAseq dataset we applied stringent filters to minimize the classification of RT-PCR artifacts as novel variants: We only considered transcript isoforms that were supported by at least two UMIs with a minimum of five reads per UMI. We eliminated transcript isoforms where exons are not joined at canonical splice sites since most false exon-exon junctions generated by RT template switching are not joined at sequences consistent with splicing by the major or minor spliceosome. We also did not consider mono-exonic transcripts since false priming on A-rich intronic regions of pre-mRNA results mainly in mono-exonic transcript isoforms (intronic priming). We also required all exon-exon junctions to be found in an E18 mouse brain Illumina short read dataset (see Methods section for details). Applying those filters, we identified 18,439 known Gencode transcript isoforms (transcripts that have all exon-exon junctions of the Gencode reference) for 11,417 genes and 15,317 novel isoforms affecting 5271 genes in our long read single-cell data.

While long read scRNAseq is well suited to analyze the diversity of transcript isoforms in single cells, other emerging techniques that do not use RT and PCR are in our opinion betier suited when the principal goal is to extend reference sequence databases such as Gencode. For instance, RNA can be sequenced directly on Oxford Nanopore sequencers, although not at a single cell level. If an oligonucleotide that serves as a 5’ tag for full-length sequences is selectively ligated to capped RNA prior to sequencing, full-length transcripts can be identified and reliable information on the diversity of transcript isoforms expressed in a tissue can be obtained^15^.

## Supplementary Data

**Supplementary Dataset 1:** E18.mouse.brain.isoforms.gff.gz,GFF file with the Gencode v18 annotated transcripts isoforms and novel transcript isoforms we identified.

